# Higher basal tumor immune activity coupled with a systemic immune response improves liver cancer outcomes

**DOI:** 10.1101/2022.04.16.488543

**Authors:** Ghazal Alipour Talesh, Daniele Vitale, Peter Langfelder, Mahmoud Karimi Azardaryany, Michael Berry, T. Grant Belgard, Jocelyn Choo, Geraint Rogers, Vikki Ho, Mehdi Ramezani-Moghadam, Suat Dervish, Joey Lai, Brian S. Gloss, Catherine L. Worth, Duncan McLeod, Mohammed Eslam, Christopher Liddle, Liang Qiao, Jacob George, Saeed Esmaili

**Author notes:** Equal authors (Ghazal Alipour Talesh, Daniele Vitale). **Correspondence and senior authors:**. **Conflicts of interest:** All authors disclose no conflicts.

## Abstract

A holistic understanding of anti-tumor immunity requires examination of systemic immunity beyond the tumor microenvironment. This link between systemic and tumor immune activity is underexplored. We demonstrate that a stronger Type I interferon response in human liver tumors predicts better survival and correlates with the immune response in the adjacent non-tumor liver. Further, in patients with liver cancer, clonal hematopoiesis (CH) (a marker of systemic immune activation) is associated with a trend towards improved survival. In a mouse model of liver cancer, basal liver immune responses correlate with the degree of bone marrow hematopoietic stem and progenitor cell responses. A higher systemic and liver immune response, marked by innate myeloid cell infiltration reduces tumor burden, while sub-optimal systemic and liver immunity corresponded with a higher tumor burden. Our findings indicate that the state of bone marrow hematopoiesis impacts liver tumor outcomes, offering both therapeutic and prognostic opportunities.

## Introduction

Environmental factors, notably hepatotropic viruses (Hepatitis B and C) are a major cause of liver cancer globally, however metabolic fatty liver disease is expected to become the dominant etiology for liver cancer within two decades [1]. These diseases modulate both the hepatic and systemic milieu and tumor microenvironment, including the composition of immune cells within the tumor. While the impact of environmental factors on the liver microenvironment and on tumorigenesis is documented [2], less understood is their effect on the systemic immune milieu and tumor outcomes. We posit that modelling these interactions can unravel novel mechanisms of tumor immune surveillance.

In humans, a robust immune response is protective against cancer and improves overall survival [3]. Conversely, an immunosuppressive tumor microenvironment is associated with poor outcomes [3]. Plausibly the interaction between tumor, host, and environmental factors determines the strength of basal immune activity within tumors. This basal immune activity state in cancer can be conceptualized as a “cancer–immune set point”, defined as the threshold over which the immune system can exert anti-tumorigenic effects. In light of this, a set point above the threshold would generate immune activity that confers a protective effect [4]. To determine the baseline immune set-point, Kotliarov *et. al.*,[5] assessed blood transcriptional signatures and gene co-expression modules. They reported that a Type I interferon (IFN) response module provides a quantitative measure of the baseline immune set point. A higher Type 1 IFN predicted improved responses to vaccination and was associated with a higher risk of flares in systemic lupus erythematosus [5]. Here, we applied similar methodologies to quantify immune activity within the liver and in tumors using gene co-expression modules.

Recent advancements in cancer profiling and modelling have shifted the focus from a tumor-cell centric perspective to one that prioritizes the tumor microenvironment. Nevertheless, more studies are needed to develop holistic, organism-wide approaches to counter cancer. Recent findings confirm the importance of non-tumor tissues and systemic immune processes on tumor outcomes[6]. A fledgling area of research is the cross-talk between bone marrow hematopoiesis, systemic immunity, and cancer development. For instance, following myocardial infarction, the expansion of Ly6C^hi^ monocytes in mouse bone marrow accelerates breast cancer progression [7]. Conversely, the induction of bone marrow granulopoiesis by β-glucan treatment elevates Type 1 IFN responses and impairs melanoma and lung adenocarcinoma growth [8]. Likewise, administering HDL nanobiologics to stimulate bone marrow myelopoiesis in mice reduces melanoma tumor growth and potentiates immune checkpoint inhibitor responses [9]. These studies suggest that the activity of the bone marrow hematopoietic system can influence tumor outcomes. Teleologically, hematopoietic stem and progenitor cell (HSPCs) responses could generate adequate numbers of adaptive and innate immune cells to sustain systemic immunity above the protective threshold to counter tumor growth. In contrast, a suboptimal HSPCs response should lead to an immune response below the threshold required to confer protection against tumors in peripheral organs. This concept is the focus of our investigations.

We first investigated bone marrow hematopoietic stem cells status and the presence of clonal hematopoiesis (CH) in patients with liver cancer. Notably, the biological effects of CH are related to hematopoietic stem cells and progenitor cells (HSPCs) [10]. In TCGA dataset, liver cancer patients with CH show a trend to better survival, suggesting that HSPCs activity could influence tumor outcomes. Subsequently we used a diethylnitrosamine (DEN) preclinical model to study the impact of bone marrow (BM) HSPCs activity on tumor outcomes. We simulated varying degrees of HSPC responses through dietary challenge. We then utilized weighted gene co-expression network analysis to assess basal liver immune activity. A heightened liver innate immune response and stimulation of HSPCs led to improved outcomes, whereas low-grade bone marrow HSPC activity increased tumor burden. This work paves the way for developing new therapeutic opportunities and holistic approaches for treating liver cancer

## Results

### A Type I IFN response in liver cancer predicts outcome and is correlated with adjacent liver tissue immune responses

We first investigated the correlation between basal liver and tumor immune activity in the TCGA dataset. These samples included paired tumor and non-tumor tissues (n=48) for which we had weighted gene co-expression network analysis (WGCNA) data available for <u>h</u>uman liver <u>c</u>ancer <u>m</u>odule *1* (hcM1) related to the immune response, and hcM54 related to Herpes simplex infection and the type I interferon response [11]. The expression level of hcM54 within tumors was associated with improved 5-year survival (hcM54, p=0.02), while higher hcM1 expression showed a trend to better survival (hcM1, p=0.06) (Figure 1A). Compared to adjacent non-tumor liver, the expression of hcM1 and hcM54 was lower in the cancer tissue (Figure 1B-C). There was a positive correlation (R2= 0.37, p= 4.510e-006) in the expression level of hcM54 (but not hcM1) between the liver cancer and the adjacent non-tumor liver (Figure 1D-E). When we examined for association *between* the expression modules (hcM54 and hcM1), a positive correlation was found both in the adjacent non-tumor liver (R2= 0.25, p= 2.462e-004) and in the tumor (R2= 0.20, p= 9.712e-019) (Figure 1F-G). Overall, these data suggests that a higher Type I IFN response in tumor tissue is predictive of better outcomes. This response was correlated with the immune response in adjacent non-tumor liver tissue. Given that a type I IFN response could represent a cancer-immune setpoint, we explored whether systemic factors and BM HSPCs activity regulates this response.

**Figure 1.**
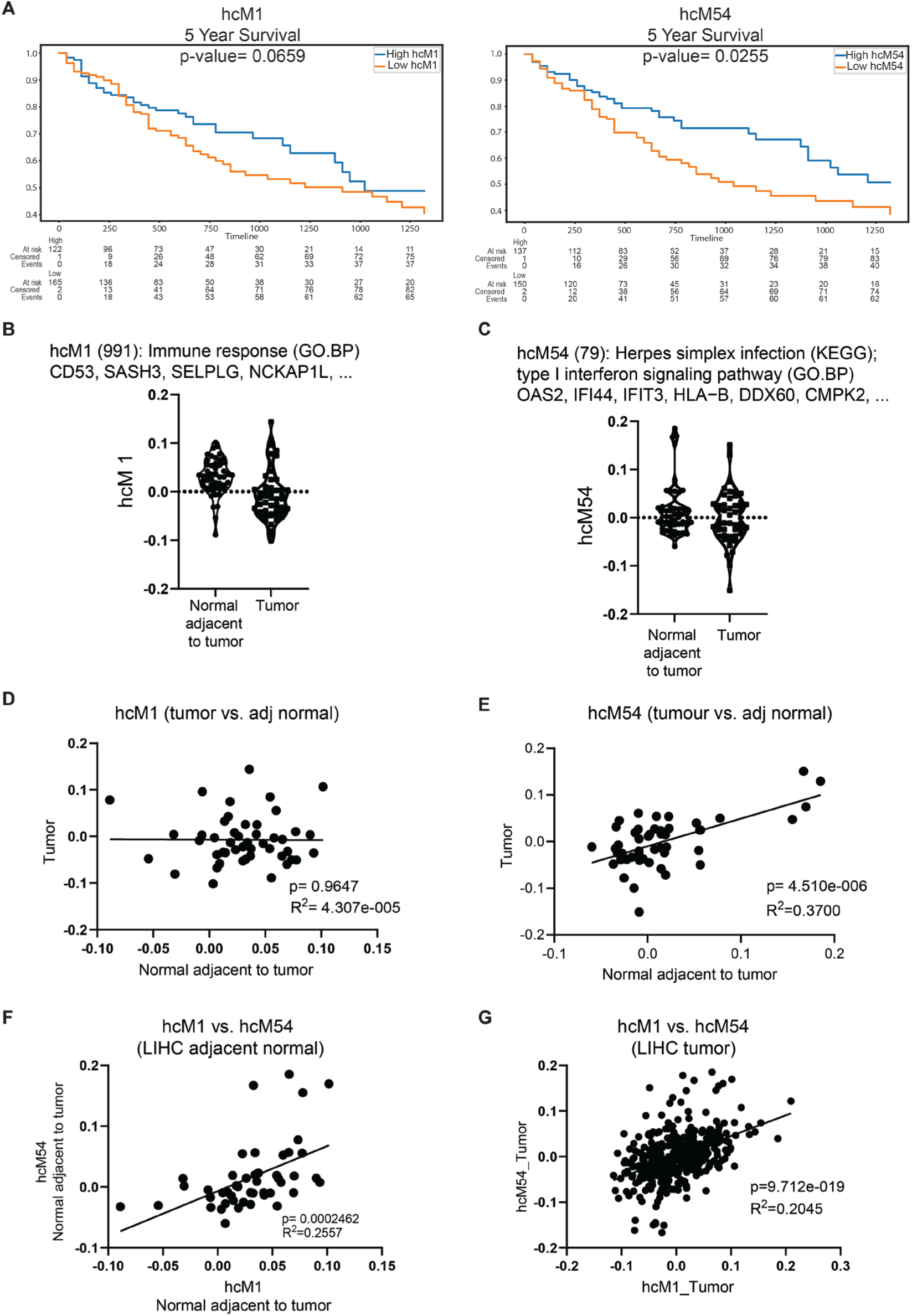
Association between the expression level of immune and interferon response modules within liver and tumors. A) Kaplan-Meier plots show that the liver tumor immune and interferon module response predicts outcomes in TCGA liver cancer dataset. B-C) Expression levels of hcM1 (immune response) and hcM54 (type I interferon response) modules in tumors and their paired adjacent non-tumor tissues in TCGA dataset (n= 48). The expression of hcM1 and hcM54 modules were lower in liver tumors. D-E) The expression level of hcM54 within tumors correlated with hcM54 levels in the paired non-tumor liver. F) There is a positive correlation between expression level of hcM1 and hcM54 within adjacent non-tumor tissues. G) The positive correlation between expression level of hcM1 and hcM54 within liver tumor tissues.

### Liver cancer patients with clonal hematopoiesis (CH) show a trend to better survival

To determine the impact of BM HSPCs status on tumor outcomes, we examined for the presence of clonal hematopoiesis which is known to be associated with liver inflammatory responses [12]. Notably, exposure to factors such as diet composition and smoking stimulate HSPCs proliferation and the earlier occurrence of CH [13]. The latter indicates elevated and sustained hematopoietic activity with higher production rates of immune cells in the marrow. We therefore explored the link between CH and tumor outcomes in the TCGA liver cancer dataset. CH was defined based on the presence of mutations in 15 well-known CH driver genes [14]. Subsequently we examined the relationship between CH and survival in TCGA human liver cancer dataset. Kaplan-Meier plots indicated that patients with CH had a trend (p=0.057) towards greater 5-year survival (Figure 2A). The available non-tumor liver samples from patients with CH (n=2) in this dataset was insufficient to investigate associations between CH and non-tumor liver immune modules (Figure 2B). This trend towards better survival suggests that higher HSPCs activity plays a role in tumor outcomes. To determine if this is true, we simulated varying degrees of HSPC responses and systemic immune activity in a preclinical model of liver cancer.

**Figure 2.**
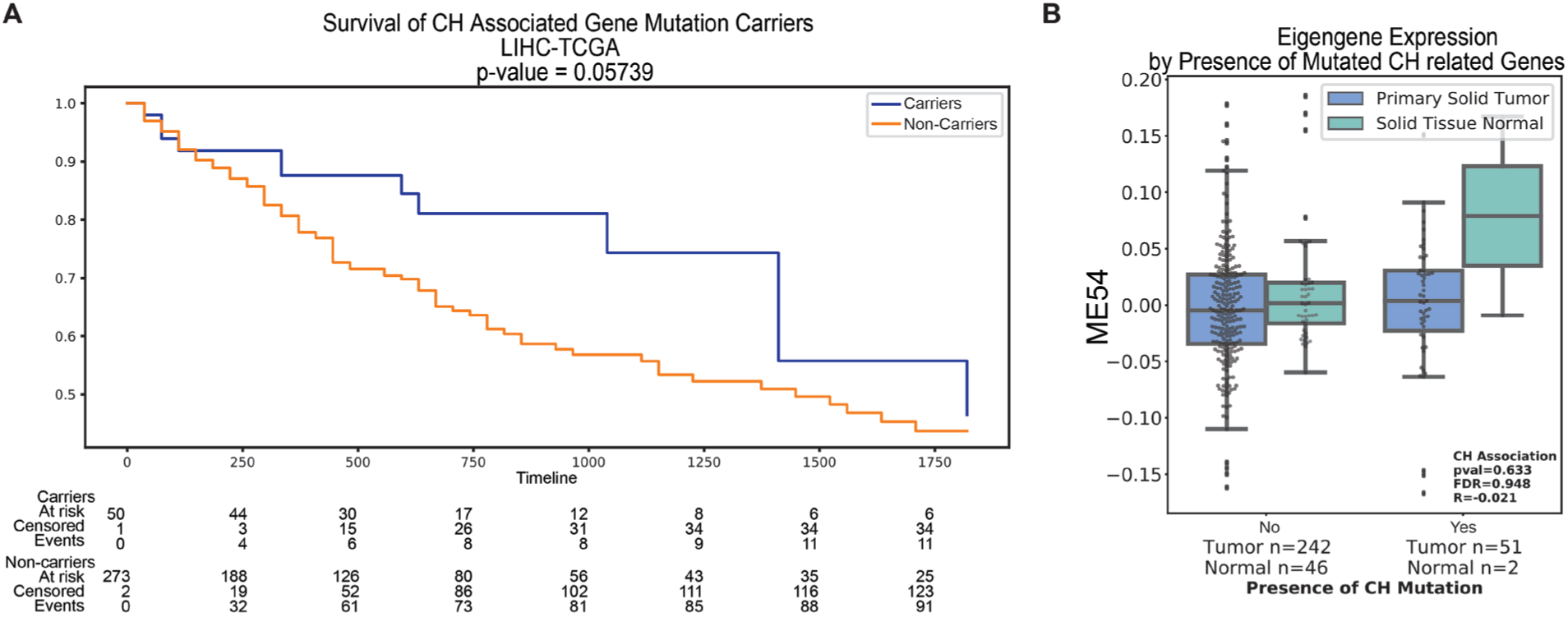
The presence of clonal hematopoiesis (CH) in liver cancer patients showed a trend to association with outcomes. A) Patients with CH show a trend (p=0.057) to better 5-years survival in TCGA. B) The number of non-tumor samples (n=2) with CH was not sufficient to explore the link between CH and liver interferon responses.

### Higher systemic immune responses confer improved tumor outcomes

To investigate the role of systemic immune responses on tumor outcomes we utilized a diethylnitrosamine (DEN) mouse model of liver cancer, and diets of varying composition to elicit differing levels of systemic immune activity. We used diets containing sucrose, cholesterol and cholic acid (HS_Chol2%_CA), hereafter referred to as a cholesterol rich diet (ChR) as dietary cholesterol is known to stimulate BM HSPCs [13]. Adding cholic acid to the diet inhibits bile acid biosynthesis from cholesterol and cholesterol secretion to the gut. Consequently, when combined with dietary cholesterol, this leads to cholesterol accumulation in the liver [15]. Additionally, we used a high sucrose (HS) diet, and a diet with added cholic acid (CA). These two diets minimally impact HSPC responses [15]. A normal chow (NC) diet was used as control (Figure 3A).

**Figure 3.**
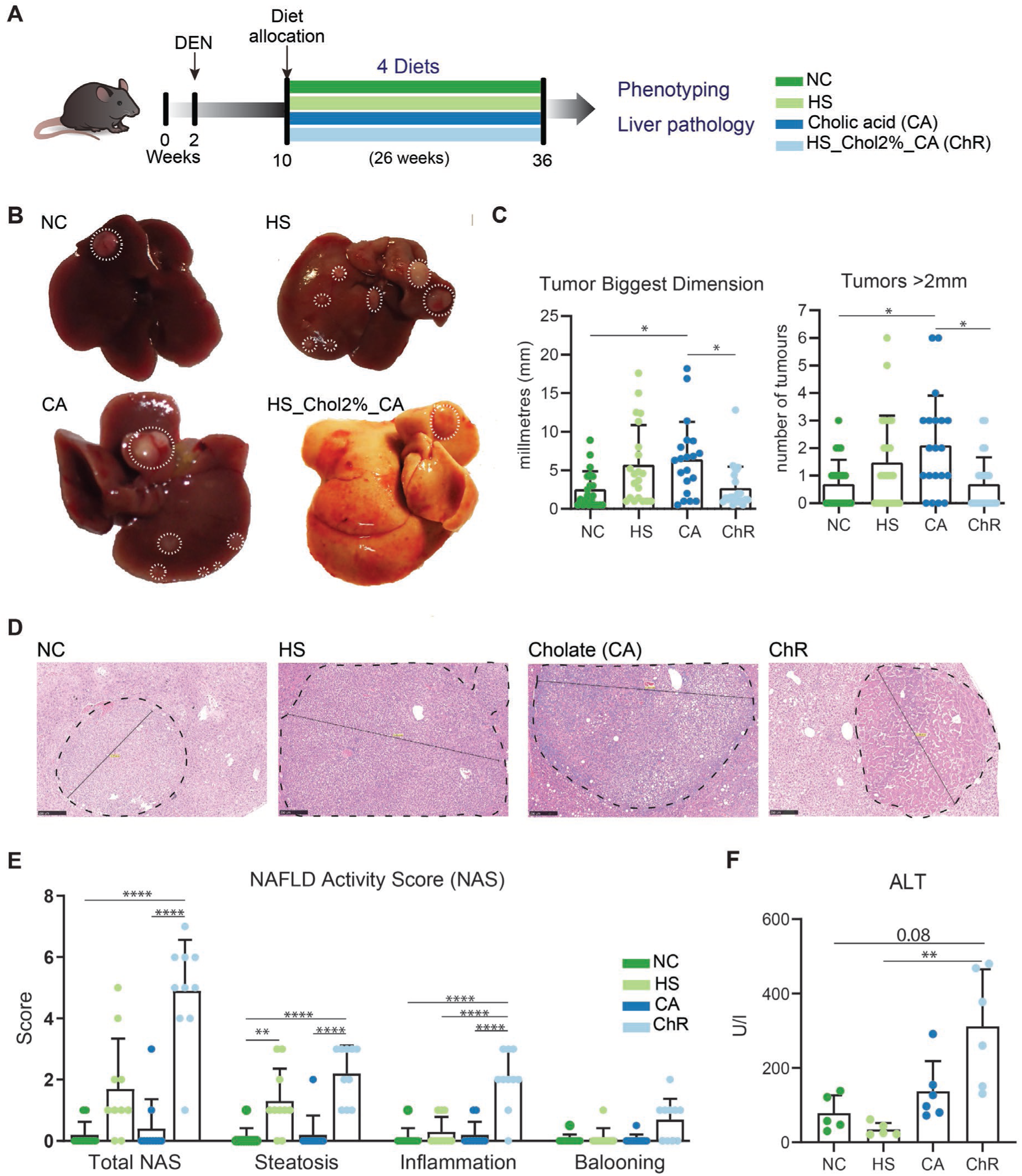
Phenotypic outcomes in a liver tumor models following dietary challenge. A) Protocol of mouse DEN tumor study. Diethylnitrosamine (DEN) (25 mg/kg) was administered at the age of 2 weeks. At 10 weeks of age mice were allocated to one of four dietary groups: i) normal chow (NC); ii) high sucrose (HS), iii) a diet with added cholic acid (CA), or iv) a cholesterol rich diet (ChR) for 26 weeks (n= 20-24 mice/group). B) Autopsy photos representing the tumor bearing livers of DEN-treated mice. C) Tumor biggest dimension and multiplicity >2mm (number of tumors with maximum diameter above 2mm per mouse). D) Representative H&E stained liver sections of DEN-injected mice fed the diets as above. E) NAFLD activity score (NAS) of non-tumor area indicates higher hepatic steatosis in HS diet fed mice with steatosis, lobular inflammation and ballooning in the ChR diet (n=10 mice/group). F) ALT levels are higher in the cholesterol rich diet group (n=5-6 mice/group was tested). Error bars represent mean ±1 standard deviation (SD). One-way ANOVA was used to test for significant differences in means with a Dunn’s (non-parametric) multiple comparison post-hoc test (Figs.1C). Two-way ANOVA with a Tukey’s (parametric) multiple comparison post-hoc test was used in Figs.1E. ****, ***, ** and * indicate P <0.0001, P<0.001, P<0.01 and P<0.05 respectively.

14-day old male C57BL6 mice were injected with the pro-carcinogen diethylnitrosamine. Starting at 10 weeks of age, mice were assigned to one of four diets for 26 weeks (Figure 3A). All mice, regardless of diet composition developed macroscopically visible tumors at 36 weeks. The CA diet group developed more and larger tumors (p <0.05) than NC and ChR groups, while a trend was also detected towards more and larger tumors in the HS group (Figure 3B-C). In contrast, mice fed the ChR diet, despite exposure to sucrose and cholic acid, had a similar tumor burden as the NC group. On histological examination, the HS group demonstrated steatosis (P<0.01), while the ChR group showed steatosis and inflammation in the non-tumor liver. We did not observe steatosis or inflammation in the non-tumor liver tissues of mice fed the CA diet (Figure 3D-E).

Before exploring the underlying mechanisms influencing tumor burden in the ChR group, we compared the anthropometric and metabolic phenotypes of the mice. Notably, the body weights of mice fed the ChR and CA diets initially declined then remained steady through the experiment (Supplementary Figure 1A). After 10 weeks on diet (week 20), glucose tolerance was impaired in mice fed the HS diet (Supplementary Figure 1B). At the end of experiment (week 36), mice fed the HS diet had markedly greater body weights, epididymal and subcutaneous fat, and liver weights (Supplementary Figure 1C-E). The ChR and CA groups had lower body weights than the control diet fed mice (Supplementary Figure 1C), but higher liver and liver/body weight ratios (Supplementary Figure 1F-G). Blood glucose level was higher in HS group (Supplementary Figure 1H). Higher triglyceride levels were observed in HS and CA groups (Supplementary Figure 1I). Further, higher blood cholesterol levels were noted in the HS and ChR groups, with highest levels in the ChR group (Supplementary Figure 1J). There was a trend towards higher ALT levels with the CA and ChR diets and to lower ALT in the HS group, supporting the presence of a high liver inflammatory response in the ChR diet group (Figure 1F). In summary, only the HS group displayed the hallmark features of metabolic syndrome. There were no significant differences in body weight or glucose homeostasis between the NC, CA, and ChR groups. Therefore, these factors cannot account for the differences in tumor burden between the CA and ChR diet groups. Despite a higher tumor burden in the HS and CA diets groups, the ChR diet group had a similar tumor burden as NC, though containing both HS and CA.

We next profiled the bone marrow hematopoietic compartment using flow cytometry (Figure 4A). In this analysis, a significant increase in the numbers of Lin-Sca1+ cKit+ (LSK) cells, short-term and long-term HSCs (HSCs), and multipotent progenitor cells (MPP2/3 and MPP4) were observed in the ChR diet fed mice (Figure 4B-D). MPP2/3 cells are myeloid biased and give rise to all myeloid linages, while MPP4 cells are biased toward lymphoid cells [16]. Interestingly, while the number of HSCs increased in the ChR diet fed mice, their frequency decreased. This phenomenon occurs when the majority of LSKs (parent cells to calculate HSC frequencies) consist of multipotent progenitor cells. Thus, the ChR diet activates BM HSCs, prompting their differentiation to progenitor cells to mount a systemic immune response. (Figure 4B-D). The CA diet also induced changes in hematopoietic stem cells but to a lesser extent. We did not detect statistically significant changes in the number and frequencies of LSK and MPP2/3 cells in the HS diet group (Figure 4B-C). In sum we demonstrate that the highest HSPCs response occurs in the ChR diet group which could play a role in modulating the tumor burden. Plausibly, the response in CA group did not reach the threshold to protect against tumor growth. Further, the minimal HSPCs response in mice fed the HS diet could not prevent tumor growth. Next, we explored whether the HSPC response is predictive of immune activity within liver.

**Figure 4.**
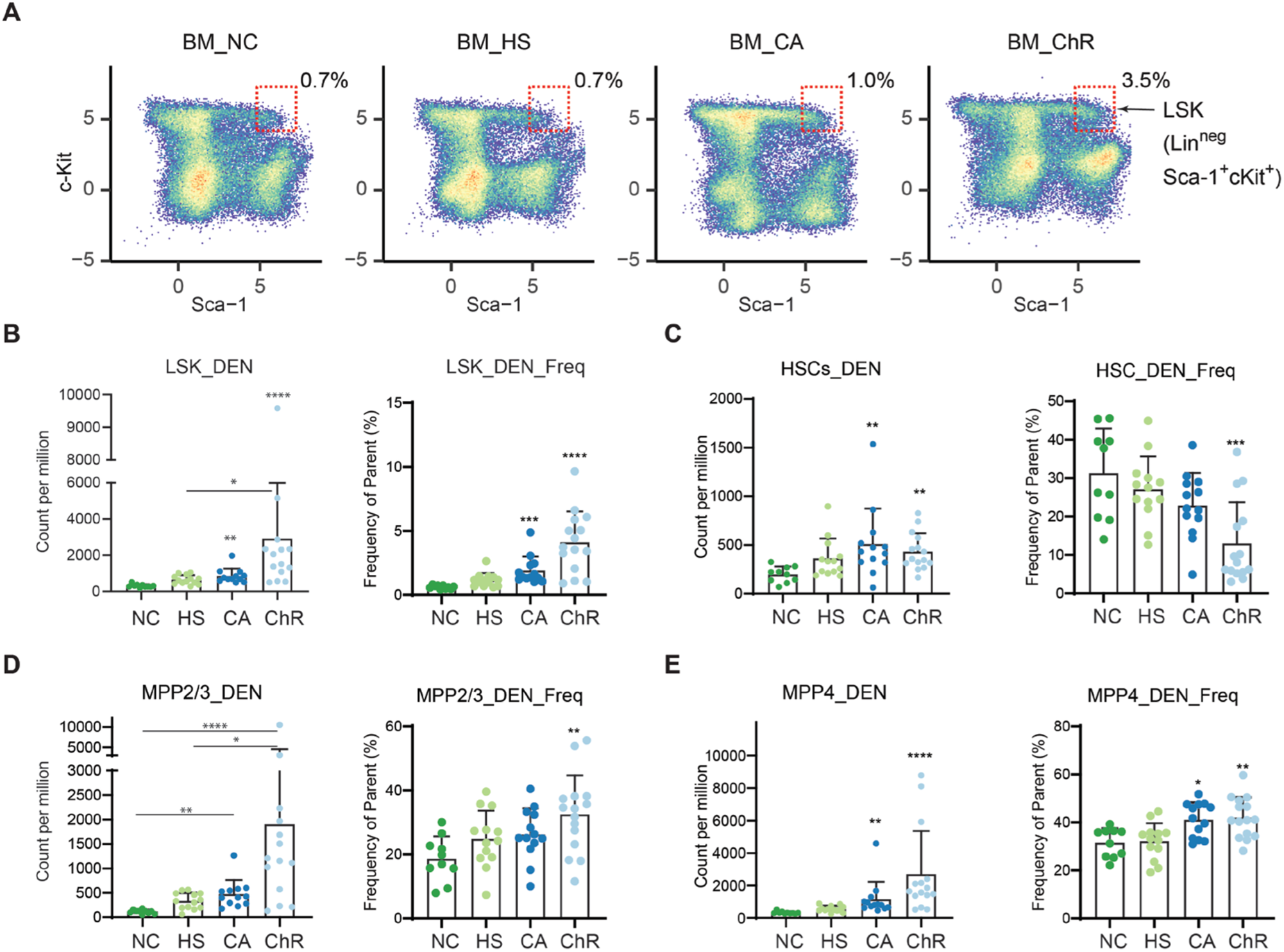
Analyzing the bone marrow hematopoietic stem cells (HPSCs) response in a mouse liver tumor model with dietary challenge. A) Representative flow cytometry plots for bone marrow LSK cells show the highest HSPCs response in the cholesterol rich diet group. B-E) Higher number of Lin-Sca-1+ cKit+ (LSK), HSCs, multipotent progenitor cell 2/3 (MMP2/3), and MPP4 in response to diets. An incremental increase in HSPCs responses from HS to CA groups, with distinct upregulation in the ChR diet fed mice. (n(_NC)_= 10, n_(HS)_= 13, n_(CA)_= 13, n_(ChR)_= 15. Error bars represent mean ±1 standard deviation (SD). One-way ANOVA was used to test for significant differences in means with a Dunn’s (non-parametric) multiple comparison post-hoc test (Figs.6C). ****, ***, ** and * indicates a significant difference compared to normal chow, with P <0.0001, P<0.001, P<0.01 and P<0.05 respectively.

To do this, we performed RNA-Seq on tumor and non-tumor liver from mice on the different diets. Principal component analysis (PCA) of non-tumor liver transcriptomes and liver tumors revealed clustering of NC and HS samples, while ChR and CA diet groups clustered together (Figure 5A-B). To better assess the extent of differences between dietary groups, we compared the genome-wide effects of diet compositions on non-tumor liver and tumors. To do this, we first permutated each diet group into three sub-groups and performed differential expression analysis and correlation analysis between non-shared subsets. This was done to prevent potential bias that could arise when a subset of samples is shared across different comparisons (e.g., NC samples are used in both ChR vs. NC and CA vs. NC comparisons). The averages of the correlations are shown in Figure 5C-D. The correlations for the ChR vs. NC, and CA vs. NC in non-tumor liver and tumor samples are 0. 68 and 0.45, respectively. This data confirms that, at the level of gene expression, CA non-tumor livers resemble ChR non-tumor liver gene expression, while the gene expression profiles in tumors are less similar.

**Figure 5.**
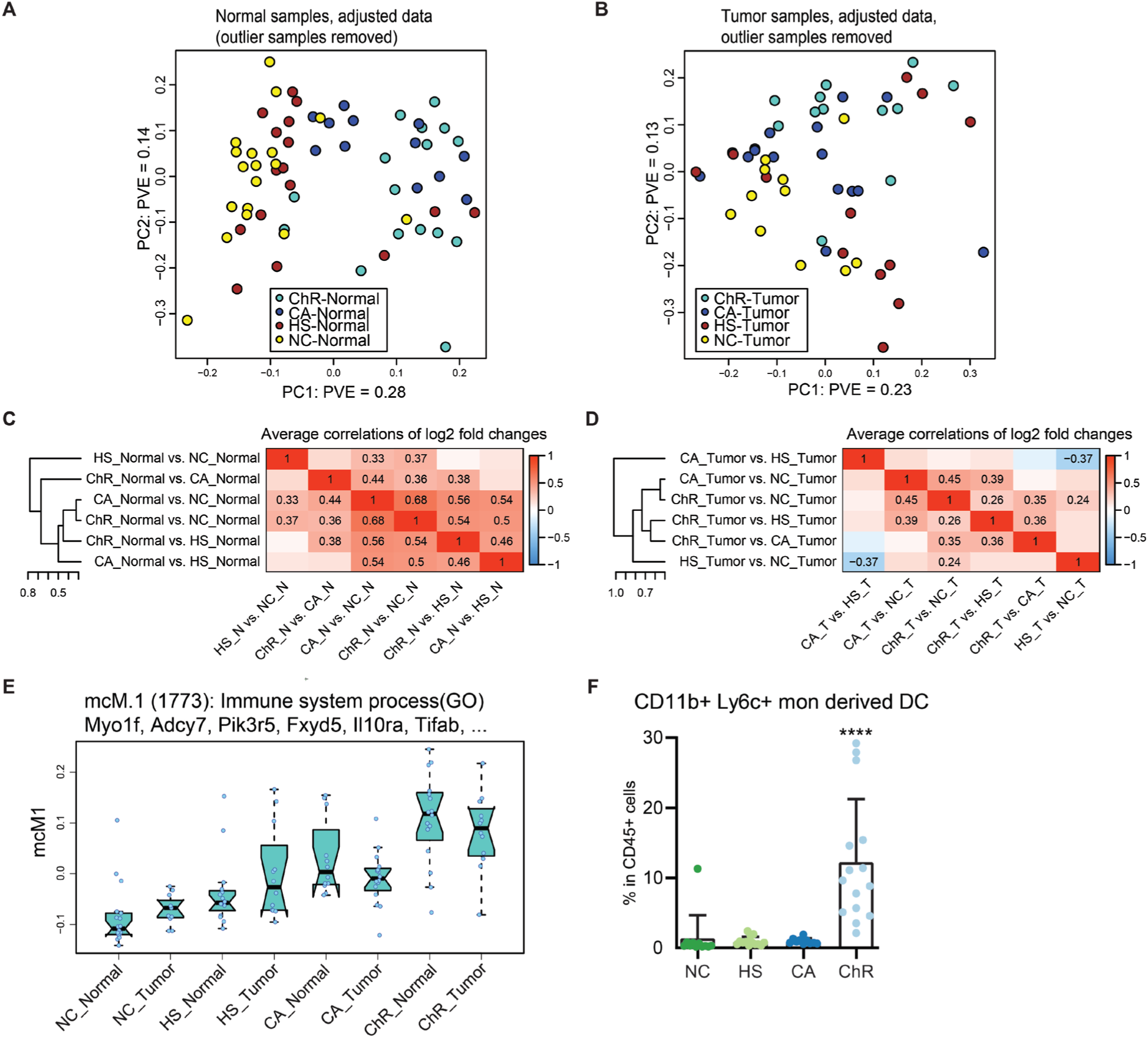
Genome wide effects of diet and tumor formation on tumor and non-tumor liver transcriptomes. A) B) Principal component (PC) plot showing the 2 leading PCs after outlier _(CA_Normal (n=2),HS_Tumor (n=1))_ removal. Each point represents one sample, color indicates diet. C-D) Genome-wide correlations of log2 fold changes for all diet contrasts in non-tumor (normal) and tumor tissues. E) Module eigengene “expression” of mouse cancer module M1 (mcM1) across the diet/condition groups. Each box represents the inter-quartile range and the thick line in the box represents the median. Whiskers extend to the most extreme point that is no further than 1.5 times the interquartile range. Blue dots represent actual eigengene expression values for each sample. The title shows the enrichment label for the module (top enrichment terms) and the top hub genes (ordered by decreasing kME). F) Immune (inflammation) scoring of H&E liver samples shows (n=10 mice/group was evaluated) similar trends as flow cytometry analysis of liver tissues. Error bars (6F-H) represent mean ±1 standard deviation (SD). One-way ANOVA was used to test significant differences in means with Dunn’s (non-parametric) multiple comparison post-hoc test. ****, ***, ** and * indicate P <0.0001, P<0.001, P<0.01 and P<0.05 respectively. Number of biological replicate used in RNA-seq: n_(NC_Normal)_= 20; n_(NC_Tumor)_= 11; n_(HS_Normal)_= 16; n_(HS_Tumor)_= 14; n_(CA_Normal)_= 14; n_(CA_Tumor)_= 16; n_(ChR_Normal)_= 15, n_(ChR_Tumor)_= 11.

Subsequently, we performed a weighted gene co-expression network analysis (WGCNA) [17] on variance stabilized data to reduce thousands of genes to a small number of transcriptionally coherent modules that represent distinct transcriptional responses to diet and disease (tumor/ non-tumor liver) (Supplementary Figure 2). Each module consists of a group of co-expressed genes and can be represented by a single representative expression profile called a module eigengene. The module preservation analysis between mouse and human liver tumors found the mouse cancer (mc) immune related module mcM1 (hub genes: *Myo1f*, *Adcy7*, *Pik3r5*, *Fxyd5*, *Il10ra*, …) as the highest preserved module (tumor Z score = 83.9) within the TCGA liver cancer dataset (Supplementary Figure 3A). The genes in mcM1 highly overlap with hcM1 (immune response) and hcM54 (Type I Interferon response) (Supplementary Figure 3B), and therefore could represent the function of these human immune related modules in mice.

Consistent with the findings in human liver tumors, the liver tumor immune response in mice follows a similar pattern as in mouse liver (non-tumor tissue). Within tumors, the expression level of mcM1 was highest in ChR tumors (Figure 5E) and they consistently had smaller tumors. The immune response in HS and CA tumors were lower than the ChR diet and they correspondingly had larger tumors (Figure 5E). Analyzing flow-cytometry data on liver samples from the mice showed enrichment of CD11bhi Ly6c + monocyte derived DCs (moDC) within the non-tumor livers of ChR mice. The presence of moDC immune cells, and the greater immune response activity (mcM1 expression) in ChR diet fed mice is consistent with a high BM HSPCs response in this group. In sum a more robust systemic immune response likely enhances liver immune surveillance which in turn reduces tumor formation and growth.

## Discussion

The tumor ecosystem consists of a heterogenous population of cells shaped by tumor cells, host, and extrinsic (environmental) factors. These elements dynamically interact to establish the basal level of tumor immune activity. There is increasing evidence that bone marrow hematopoietic system activity influences the tumor microenvironment and outcomes. Hence, to better understand this effect on liver tumor outcomes, we examined for the presence of clonal hematopoiesis. A trend towards improved survival in CH-positive patients was observed in this small cohort from TCGA tumor dataset, pending validation in a larger cohort. To explore further, we used a preclinical liver cancer model to examine the association between different degrees of BM HSPCs activity and tumor outcomes. In this analysis, the extent of immune responses in both tumor and non-tumor liver corresponded to the degree of bone marrow hematopoietic stem and progenitor cell (HSPC) activity. A cholesterol-rich diet heightened the anti-tumor immune response, characterized by significant increases in monocyte derived DCs infiltration and a corresponding decrease in tumor burden. Conversely, diets that elicited minimal HSPCs and liver immune activity were associated with larger tumors.

Exposure to environmental factors such as smoking and high dietary cholesterol stimulates HSPCs and leads to earlier appearance and higher percentage of clonal hematopoiesis [13] and consequently, a higher production of clonal immune cells from BM HSPCs. Notably, clonal hematopoiesis has been shown to be associated with liver inflammation [12]. The role of clonal hematopoiesis in solid tumors and their impact on response to therapies is also a rapidly evolving area of research. There are reports that clonal hematopoiesis is associated with poor responses to immune checkpoint inhibitors (ICIs) [18], though this relationship appears to be context-dependent, with conflicting reports in colorectal cancer [18-19]. A recent analysis showed clonal hematopoiesis-associated immune cells within tumors drives an inflamed tumor microenvironment with poor survival outcomes, however their presence predicts a better response to ICIs [20]. Particularly TET2-driven CH is associated with transcriptomic and proteomic signatures of ICI response in colorectal adenocarcinoma, glioblastoma multiforme, and lung adenocarcinoma [20]. Of note, the discrepancies between studies may arise from grouping different types of mutations under the umbrella of clonal hematopoiesis. For instance, clonal hematopoiesis with TET2 mutations—but not clonal hematopoiesis with DNMT3A mutations—has been associated with HCC development [21]. Further, TET2-driven hematopoiesis is associated with IL-1B production and heightened IL-6 and IL-8 levels [21-22]. In contrast, environmental exposure such as infections and chronic inflammation, with elevated IFNγ levels may select for clones with *DNMT3A* mutations [23]. Future studies with larger sample sizes can discern the effects of specific genetic variants in the hematopoietic system to provide better insights. Targeting the dysfunctional or mutated clones may be the future of clone-specific hematopoietic therapies in cancer [24].

In this study we demonstrated that dietary composition regulates hematopoiesis. Inducing low-grade (with HS or CA diets) liver inflammation increased liver cancer burden. In contrast, heightened hematopoietic responses with infiltration of monocyte derived DCs, and a robust type I IFN response reduced liver tumor burden. The Type I interferon response signature, as determined by Kotliarov *et. al.*,[5] was also enriched in pDCs and moDCs. Ideally, therapeutic strategies should optimally regulate the hematopoietic and liver immune response, adapting to different clinical contexts. These strategies should control low-grade inflammation to prevent liver fibrosis progression and tumor development, but in the context of established tumors should enhance the type I IFN response.

Our work underscores the potential of quantitative measures of type I IFN response and hematopoietic system fitness for diagnostic and therapeutic purposes in chronic liver diseases and cancer. The efficacy of anti-tumor immune therapies could be ensured by maintaining a type I interferon response above the therapeutic threshold. Coupling these therapies with strategies to enhance systemic immune responses could improve patient outcomes. Finding predictive markers of local tumor and systemic immune responses would streamline decision-making processes on therapeutic planning and treatment duration.

## Supplementary figures and figure legends

**Supplementary Figure 1.**
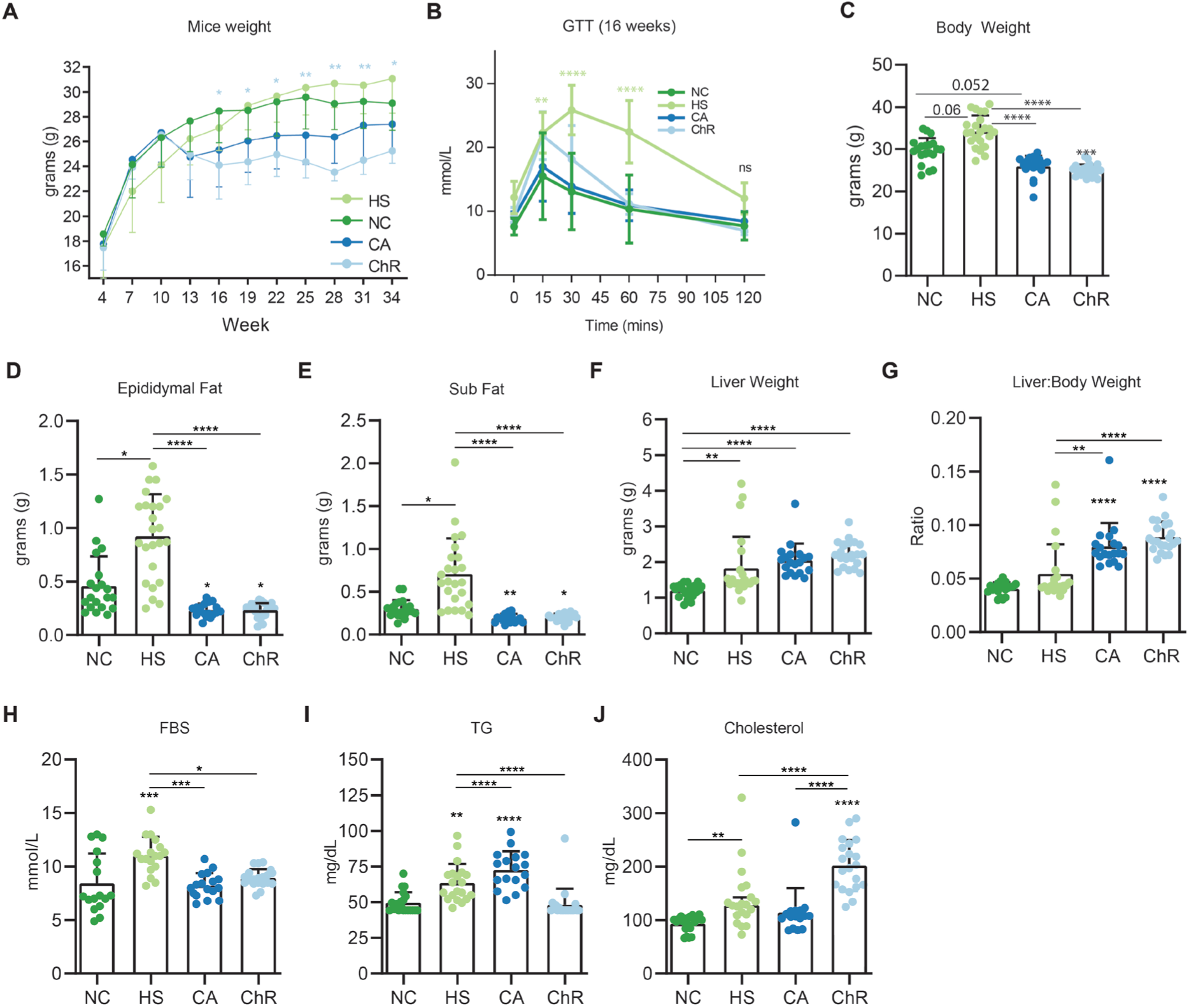
The phenotypic characteristics of mice in DEN and diet study. A) Longitudinal mouse body weights. B) Glucose tolerance test (GTT) shows impairment in glucose homeostasis in the HS group (n=6-9 mice/group was tested). C) Body weight at experiment end (36 weeks). D-E) Higher epididymal and subcutaneous fat in HS diet group. F-G) Liver weight and liver/body weight ratio at experimental end (36 weeks). H) fasting blood glucose (FBS) at 36 weeks. I-J) Serum triglyceride and cholesterol levels at week 36. Error bars represent mean ±1 standard deviation (SD). Two-way ANOVA with Tukey’s (parametric) multiple comparison post-hoc test was used in Supplementary Figure1B. One-way ANOVA was used to test for significant differences in means with Dunn’s (non-parametric) multiple comparison post-hoc test (Supplementary Figures.1C-J). ****, ***, ** and * indicate P <0.0001, P<0.001, P<0.01 and P<0.05 respectively.

**Supplementary Figure 2.**
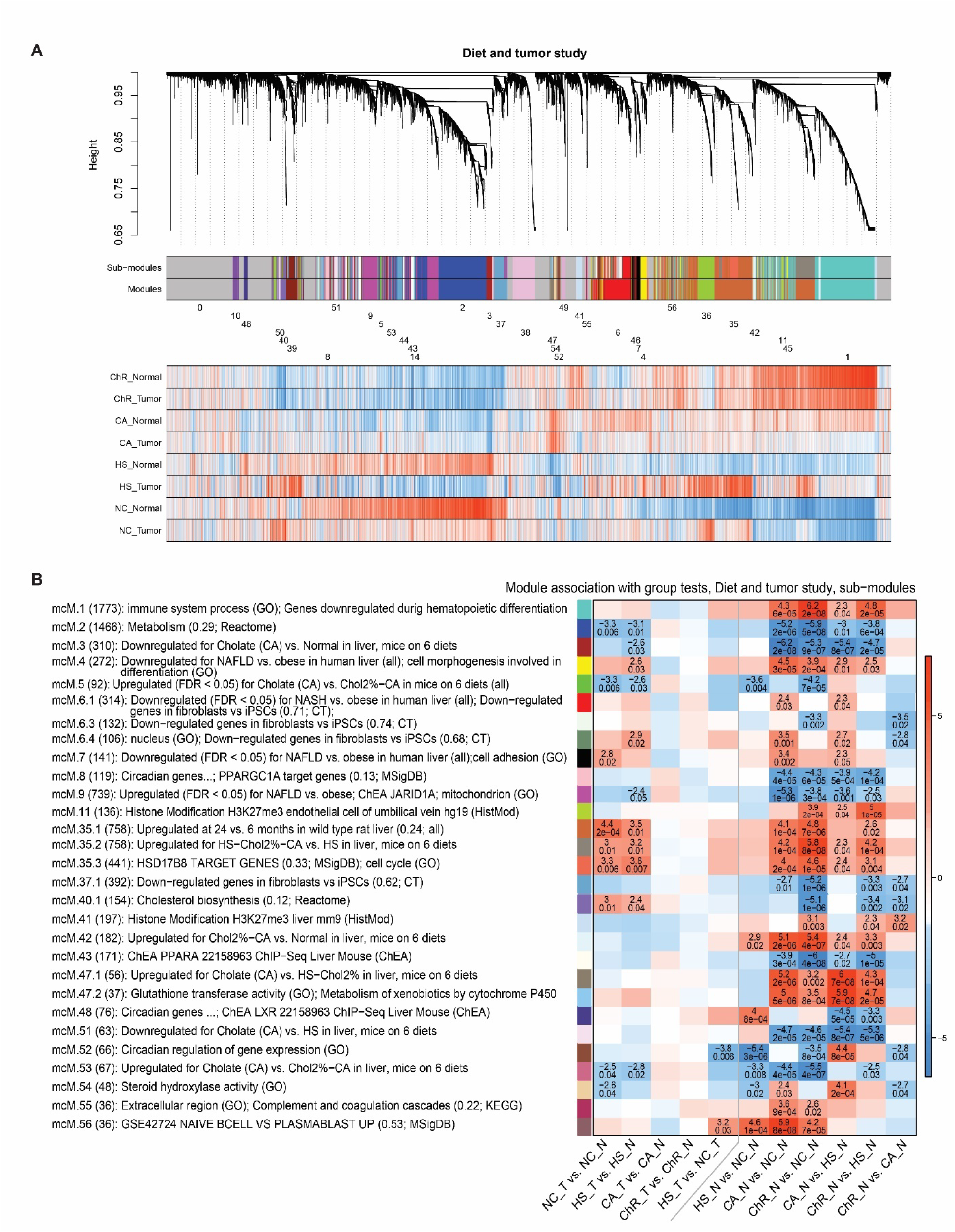
Co-expression network analysis of normal and mouse tumor liver samples. A) Gene clustering tree (dendrogram) of data from all samples (tumor and non-tumor and tumor samples) Under each clustering tree, two color rows indicate sub-modules and main modules. Numeric module labels are shown for the main modules. Below the numeric module labels, heatmaps show mean gene expression across diet and condition groups. For the heatmaps, each gene expression profile was scaled to mean 0 and variance 1. Blue and red colour indicates low and red high expression, relative to overall mean for each gene. B) Association of selected module eigengenes of sub modules across all samples with diet and condition (tumor vs. normal) contrasts. Each row corresponds to one of the modules strongly associated with at least one contrast. Row labels indicate the numeric module label, module size and top enrichment terms. Color rectangles correspond to the module colour in Figure 4A. Each column corresponds to a diet or condition contrast. In the heatmap, numbers give the association significance Z statistic and the FDR estimate. For clarity, Z and FDR are only shown for those cells where FDR<0.05. Color indicates strength and direction of the association Z statistic.

**Supplementary Figure 3.**
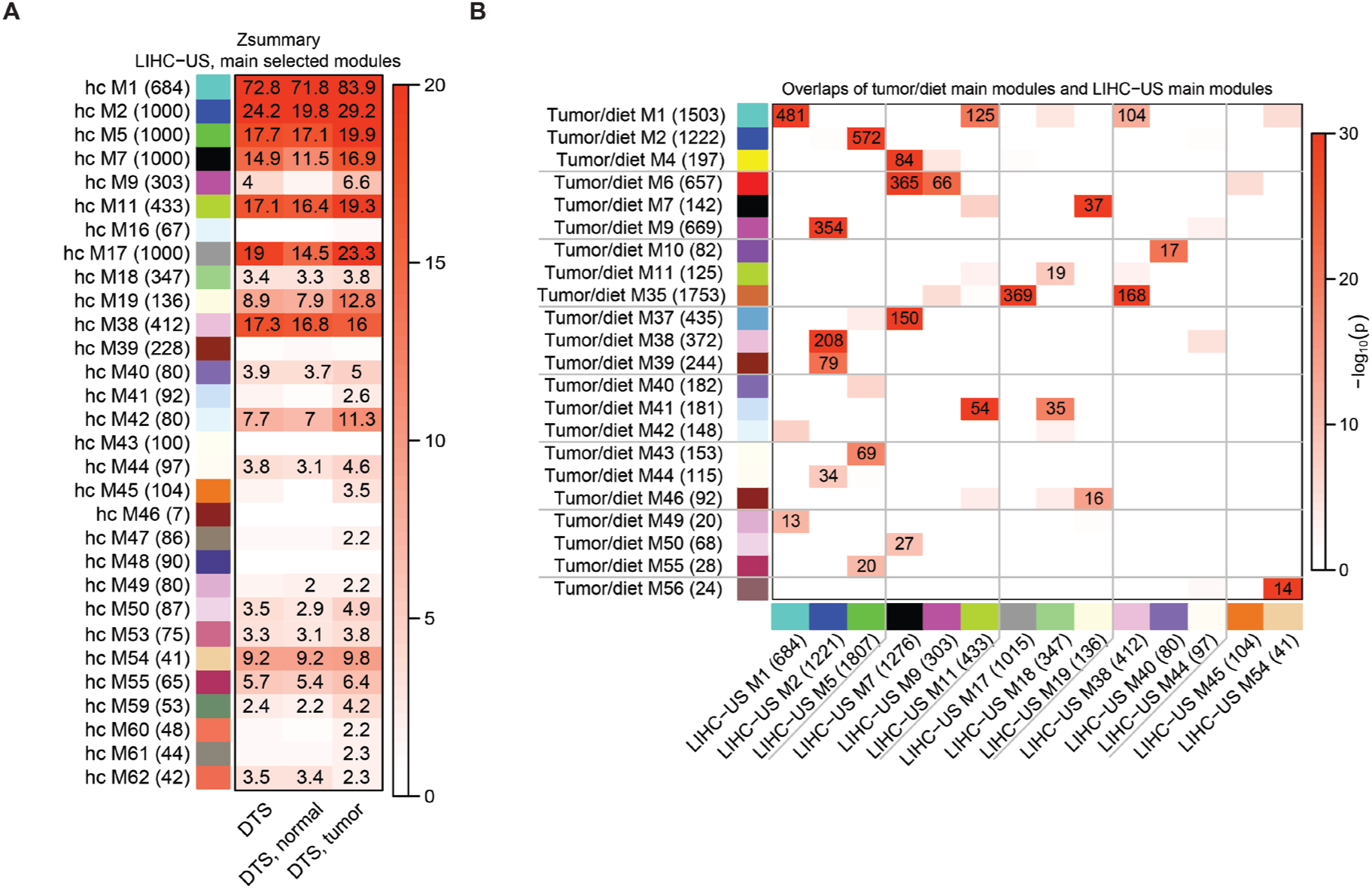
Reciprocal module preservation analysis demonstrates immune response is highly preserved between mouse and human liver cancer. A) Heatmap representation of module preservation Zsummary statistics of diet/tumor networks constructed in the DEN tumor study, and human liver cancer (TCGA). The highest preserved module is immune related module (hcM1). B) Module overlap of samples in the mouse DEN tumor study with human liver cancer.

## Experimental Section

### Clonal Hematopoiesis

The somatic mutations for LIHC samples were extracted form dbGap submission (prj33479). Low coverage variants (SNPs < 7; InDels <10) and those with an allele frequency lower than 2% or fewer than 3 alternative reads were removed from further analysis. A list of clonal hematopoesis (CH) associated genes and variants was obtained from Intogen and used to annotate LIHC samples as carriers of CH related gene mutations. To determine the association between module expression and presence/absence of CH related mutations, we performed Point Biserial correlation between ME expression levels and binary presence/absence of CH mutations.

### Mouse studies

All procedures were approved by the Western Sydney Local Health District Animal Ethics Committee and conducted in accordance with Animal Experimentation guidelines of the National Health and Medical Research Council (NHMRC) of Australia. Male C57BL/6 mice were obtained from Animal Resources Centre (Perth, Australia) and used for establishing breeding colonies at the Westmead Research holding facility and their litters were used in the diet studies. Mice were exposed to a 12-hr light/dark cycle in pathogen free conditions with free access to food and water. Male C57BL6 mice were injected intraperitoneally with 25 mg/kg body weight DEN (Sigma-Aldrich, Munich, Germany) at 14 days of age. These mice were then given either a sucrose rich diet (HS), a cholesterol rich (ChR) diet containing high sucrose with 2% cholesterol and 0.5% cholic acid (HS_Chol2%_CA), or a diet containing 0.5% cholic acid (CA) (Specialty Feed Service, Glen Forest, Australia) starting at 10 weeks of age for 26 weeks (n=20-24 mice per group). The control mice group were fed NC (Specialty Feed Service, Glen Forest, Australia) throughout the experiment. The mice were harvested at the age of 36 weeks. At the time of harvest, mice were anesthetized with i.p. ketamine (100 mg/kg)/xylazine (10 mg/kg) injection after a 4-hr fasting period. Blood was collected by cardiac puncture. Liver was harvested, and a small portion of normal liver and resectable tumors was rapidly snap frozen in liquid nitrogen and stored at -80 °C. A thin slice of liver tissue was formalin fixed for histology examination.

### Liver histology and analysis

Liver tissues were fixed in 10% neutral-buffered formalin for 24 hrs and paraffin-embedded tissue sections were stained with hematoxylin and eosin (H&E). The tissues had undergone whole tissue scanning (Hamamatsu Photonics, NanoZoomer, C9600-02). The samples were blindly assessed by pathologist DM according to the criteria for grading and staging of murine fatty liver[25] and liver tumors [26].

### Glucose tolerance test

The glucose tolerance test (GTT) was performed to evaluate mice capacity in processing and storing glucose. Fasting blood glucose (FBS) (after a 3-hour fasting period) was measured by taking blood from the tail tip using an Accu-Chek glucometer (Roche Diagnostics). Mice then received an intraperitoneal (ip) injection of glucose (1.5 mg/g), and blood glucose readings were taken at 15-, 30-, 45-, 60- and 120-mins post injection.

### Biochemical assays

Serum ALT was measured by automated techniques within the Department of Clinical Chemistry, Westmead Hospital. Serum lipids were measured using the Cobas b 101 POC system and its lipid disc, both from Roche Diagnostics.

### Flow cytometry and analysis

The mouse femurs and tibias were collected in a petri dish containing ice-cold PBS. They were cleaned by removing skin and tissue using a scalpel & forceps and the ends of the bone were carefully cut with a scalpel or razor blade. This was followed by flushing out the marrow with cold PBS (1-2 mL) using a 5 mL syringe and 30 G needle. The cells were collected in a 50 ml tube, mixed gently to make a homogenous suspension and spun for 5 min at 1500 rpm (4°C). The supernatant was gently discarded, the pellet was suspended in 5 ml 1X RBCs lysis buffer (BD Pharm Lyse™) and incubated at room temperature for 10 min. The reaction was stopped by the addition of 10 ml PBS and the tube centrifuged at 500 x g for 5 min at 4°C. The supernatant was discarded, and the pellet was suspended in 1000 µl FACs buffer (90% PBS, 5% EDTA, 5% FBS). The live singlet cells were counted using Trypan Blue in a 1:20 dilution by mixing 10μl of cell suspension with 190 μl of Trypan Blue. For bone marrow stem cells staining, 6x10^6^ cells/sample were distributed into a V bottom 96-well plate and assessed for Cell viability by incubation with 50µl Zombie yellow live dead (1:500 in PBS) for 15 min in the dark at room temperature. The cells were then topped up with 200 µl PBS, spun down at 500 x g for 5 min at 4°C. The supernatant was gently sucked off and the pellet suspended in 50 µl of antibody cocktail in Brilliant violet (BV) stain buffer for 20 min in the dark at room temperature. After 30 minutes of antibody staining, the cells were topped with 200 µl FACS buffer, spun down at 500 x g for 5 min at 4°C. The supernatant was then gently aspirated, and the pellet was washed 3 times with 200 µl FAC buffer and spun down at 500 x g for 5 min at 4°C. After the final wash, the pellet was suspended in 300 µl FACS buffer and transferred to a FACS tube and kept on ice in the dark. One well of unstained cells was also included in all experiments. Single colour controls (SCCs) were prepared by adding only one single antibody to the FACS buffer. Fluorescence minus one (FMO) controls were prepared by adding all the antibodies of each panel except for one to the FACS buffer. A pooled mixture of all samples was used for unstained wells, SCC and FMOs. Fluorometric data were acquired on a flow cytometer (Fortessa, Becton Dickinson).

Raw FCS files were compensated, arcsinh transformed (a=0, b=1/150, c=0), cleaned (margin events, debris, singlets, live) and cell populations of interest (Lin-,Sca-1+,c-Kit+ [LSK], Lin-,Sca-1-,c-Kit+ [LKS-] and residual Lin-) were gated using functions provided by ggcyto_1.18.0, openCyto_2.2.0, flowStats_4.2.0 and flowCore_2.2.0 [27]. CATALYST_1.14.0 [28] provided wrapper functions for FlowSOM and ConsensusClusterPlus. Pre-gated LSK, LKS- and residual Lin-cells were clustered based on cell marker expression (fluorescence intensity). Similar clusters were manually merged into cell subtypes of interest as defined here [29]. Plots were generated using ggplot2_3.3.3. The glm function provided by glmm_1.4.2 was used to fit linear models (family = gaussian) where cells/million live singlets of studied populations and diet were the response and independent variables, respectively. Pairwise adjusted Pvalues were generated using Tukey’s honest significance test using the glht function provided by multcomp_1.4-17. The analyses were performed in R 4.0.3 global environment using RStudio IDE 1.2.5033.

Antibodies: CD45 (BUV 395), Cat. No.: 564279, BD Biosciences; CD16/32 (BUV 737), Cat. No.: 565272 BD Biosciences; CD34 (BV 421), Cat. No.: 562608 BD Biosciences; CD127(BV 510), Cat. No.: 135033 Biolegend; Zombie Yellow, Cat. No.: 423103 Biolegend; CD150 (BV 650), Cat. No.:115931 Biolegend; Sca-1 (BV 711), Cat. No.: 108131 Biolegend; CD90.2 (BV 786), Cat. No.: 105331 Biolegend; CD29 (FITC), Cat. No.:102206 Biolegend; Lin Cocktail (PerCP-Cy5.5), Cat. No.: 561317 BD Biosciences; CD135 (PE), Cat. No.: 553842 BD Biosciences; CD140a (PE-Cf594), Cat. No.: 562775 BD Biosciences; CD105 (PE-Cy7), Cat. No.: 120410 Biolegend; CD117 (c-Kit) (APC), Cat. No.: 553356 BD Biosciences; CD48 (APC-Cy7), Cat. No.: 103432 Biolegend.

### FACS analysis on intrahepatic leucocytes

Intrahepatic leukocytes were isolated and subjected to FACS analysis as described previously [30]. Cells were stained for viability using Zombie Fixable Viability Kit for 20 mins. Prior to antibody staining Fc receptor blockade was performed with the use of TruStain fcX™, anti mouse CD16/CD32 (BioLegend Clone 93 Cat. 101319). Samples were then stained with the following antibodies: Anti CD45 (30-F11; Cat. 564279), anti CD4 (GK1.5; Cat. 564298), anti NK1.1 (PK136; Cat. 562921), anti CD3e (145-2C11; Cat. 561100) (all from BD Biosciences) and anti MHC-II (M5/114.15.2; Cat.107635), anti CD8a (53-6.7; Cat. 100747), anti Ly6C (HK1.4; Cat. 128041), anti CD11b (M1/70; Cat. 101206), anti CD11c (N418; Cat. 117347), anti F4/80 (BM8; Cat. 123116) and anti Ly6G (1A8; Cat. 127622) (all from BioLegend). Flow cytometric analysis was performed on a flow cytometer (LSR II Fortessa, Becton Dickinson) and analysed with FlowJo (Tree Star, Ashland, OR, USA).

### RNA_Seq

Total tumor and non-tumor liver RNA were extracted using RNAeasy (Qiagen). RNA purity and integrity were confirmed using Tapestation. Libraries were prepared from 100 ng total RNA using the Illumina Truseq stranded mRNA library prep kit. The library average fragment length and standard deviation (SD) was verified using Tapestation before 100bp-SE sequencing on the Illumina NovaSeq 6000, yielding a median of 20 M reads per sample for the mouse tumor study. Sequencing adapters and low-quality reads were trimmed using the ‘Maximum information quality filtering’ algorithm implemented in Trimmomatic-0.39. A cut-off length of 40 and strictness setting 0.999 was used [31]. The cut-off for the minimum read length was set to 36bp.

Reads were mapped using Salmon-1.4.0 [32] . against the GENCODE vM25 mouse transcriptome with GRCm38 mouse genome assembly as a decoy. The Salmon mapping algorithm was used with both random hexamer priming and GC bias correction.

The R package tximeta-1.8.2 was used to generate raw count matrices from Salmon quant.sf output files. Reads were summarized at the gene level and raw count matrices exported as csv for downstream processing.

## Quantification and statistical analysis

### Statistical analysis of phenotypic data

Bar and XY graphs presenting mouse phenotypic data (Figure 1 and Supplementary Figure 1) were generated in GraphPad prism version 8.4.3. Error bars represent ±1 standard deviation (SD) from the mean. The data presented are the pool of two independent experiments.

One-way ANOVA with Dunn’s (non-parametric) multiple comparison post-hoc test was used to test for significant differences in means. Two-ways ANOVA with Tukey’s test was used in Fig1B, 1H, and Supplementary Figure1A. Statistical significance is presented as *P*<0.05 (*), *P* <0.01 (**), *P*<0.001 (***), or *P*<0.0001 (****).

### WGCNA, enrichment, and preservation analysis RNA_Seq data pre-processing

We retained only genes with at least 0.5 counts per million reads in at least the number of samples in the smallest group (i.e., 12 samples). The rationale is to only include genes that are likely to be expressed in at least 1 diet/condition group.

To identify outlier samples, we used a modified version of the sample network methodology originally described by Oldham et.al. [33]. Specifically, to quantify inter-sample connectivity, we first transformed the raw counts using variance stabilization (R function varianceStabilizingTransformation) and then used Euclidean inter-sample distance based on the scaled profiles of the 8000 genes with highest mean expression. The intersample connectivities *k* were transformed to Z scores using robust standardization,

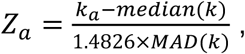

where index *a* labels samples, MAD is the median absolute deviation, a robust analog of standard deviation, and the constant 1.4826 ensures asymptotic consistency (approximate equality of MAD and standard deviation for large, normally distributed samples). Finally, samples with *Z_a_* < -6 were removed. This procedure resulted in the removal of 2 normal (CA diet) and 1 tumor (HS diet) sample.

To make DE testing and WGCNA analysis robust against potential outlier measurements (counts) that may remain even after outlier sample removal, we calculated individual observation weights designed to downweigh potential outliers as follows. First, separately for each gene, Tukey bi-square-like weights *λ* [34] were calculated for each (variance-stabilized) observation *x_a_* (index *a* labels samples) as

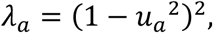

where

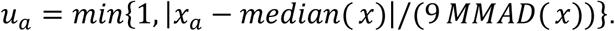

The median was calculated separately for each gene across all samples. MMAD stands for modified MAD, calculated as follows. For each gene, we first set MMAD = MAD. The following conditions were then checked separately for each gene: (1) 10^th^ percentile of the weights *λ* is at least 0.1 (that is, the proportion of observations with weights <0.1 is less than 10%) [35] and (2) for each individual group (diet), 40^th^ percentile of the weights *λ* is at least 0.9 (that is, at least 40% of the observations have a high coefficient of at least 0.9). If both conditions are met, MMAD = MAD. If either condition is not met, MMAD equals the lowest value for which both conditions are met. The rationale is to exclude outliers but ensure that the number of outliers is not too large either overall or in each genotype group. This approach has previously been used in [36],[37].

### Differential expression analysis

DE testing was carried out in R using package DESeq2 [38] version 1.28.1, with each diet/group contrast as a variable of interest. We disabled replacement of outlier measurements in DESeq2 (because outliers are suppressed using weights) as well as independent filtering of the results (because low-expressed genes were filtered out in the preprocessing step.

### Enrichment of sets of DE genes

We evaluated enrichment of significantly DE genes in the following collections of literature sets:

1. Gene Ontology (GO)[39];
2. Molecular Signatures Database (MSigDB) version 6.2 [40];
3. NCBI BioSystems gene sets[41], including KEGG, Reactome, Lipid Pathways and BioCYC;
4. Cell type markers for various immune and blood cells;
5. Liver-related sets from a collection of gene sets curated from general literature;
6. Genomic position gene sets; each set contains genes in a 5 Mb window, with two adjacent windows overlapping by 2.5 Mb;
7. Enrichr [42] 2016 ChEA library;
8. Enrichr 2015 ENCODE histone modification library;
9. Enrichr 2015 ENCODE TF ChIP-seq library;
10. Enrichr 2017 mirTarBase library [43];
11. Sets of genes DE at FDR<0.05 in our previous study of pre-clinical dietary models of fatty liver disease [36];
12. Sets of genes DE at FDR<0.05 in a study of human fatty liver disease[44] .

These 12 collections contain a total of 44,483 gene sets. Down- and up-regulated gene sets were tested separately. The calculations were carried out using R package anRichment (https://horvath.genetics.ucla.edu/html/CoexpressionNetwork/GeneAnnotation/) that implements standard Fisher exact test and a multiple-testing correction across all query and refence gene sets, resulting in a stringent Bonferroni multiple testing correction.

### Weighted gene co-expression network analysis Module identification

WGCNA was carried on variance stabilized data from 3 sets of samples: all (“Diet and tumor study”), normal (“Diet and tumor study, normal samples”) and tumor samples (“Diet and tumor study, tumor samples”). Briefly, WGCNA [45] starts by constructing a Topological Overlap (TO) matrix [46]. To this end, we used weighted correlation with individual sample weights determined as described above and the “signed hybrid” network in which negatively correlated genes are considered unconnected. Based on scale-free topology analysis, we used the soft thresholding power β=6 in the analysis of all 3 data sets.

WGCNA uses topological overlap-based dissimilarity as input to average-linkage hierarchical clustering that results in a dendrogram. Modules are identified as branches in the dendrogram using Dynamic Tree Cut [47]. To improve stability of the found modules, we used random subsets of approximately 0.63 times the number of samples to create 50 sets of perturbed WGCNA module labels. These are then used as an additional input to Dynamic Tree Cut in the final module identification. We used two different settings of Dynamic Tree Cut to produce two sets of module labels, one resulting in larger, more robust modules (called the “main” modules) and the second that produces a finer split of those main modules that can be thought of as consisting of 2 or more sub-modules. Module labels for modules from all samples were chosen such that modules that strongly overlap with modules from our previous study of pre-clinical dietary models of fatty liver disease [36] carry the same label, if possible. Module labels in the normal and tumor subsets were chosen similarly but all-sample modules were used as reference.

After module identification, we analyzed the enrichment of genes in each module in the same reference collections of gene sets that we used for enrichment analysis of DE genes. To make it easier to keep track of the biological function of each module suggested by enrichment analysis, we created a label (“enrichment label”) for each module that collects top enriched terms from each of the 12 collections. The enrichment labels are constructed only from enrichment terms whose Bonferroni-corrected p-value is below 0.05 and the overlap of the term and module covers at least 10% of the module.

Module eigengenes lead to a natural measure of similarity (membership) of all individual genes to all modules. We define a continuous (“fuzzy”) measure of module membership of gene *i* in module *I* as

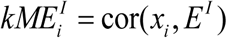

where *x_i_* is the expression profile of gene *i* and *E^I^* is the eigengene of module *I*. This definition is applicable to every individual network (data set). The value of module membership lies between -1 and 1. High *kME* indicates that the expression profile of gene *i* is similar to the summary profile of module *I*. Since we use signed networks here, we consider module membership near -1 low. The advantage of using correlation to quantify module membership is that the corresponding statistical significance (p-values) can be easily computed. Genes with highest module membership are called hub genes.

Association of modules with diet and condition was tested using linear models implemented in R package limma [48].

### Module preservation between current and previous network analyses

We carried out module preservation calculations between the current 6 network analyses and our previous study of pre-clinical dietary models of fatty liver disease [36], in human fatty liver data [44] and 3 human liver cancer data sets (LICA-FR, LIHC-US and LIRI-JP) from the ICGC portal [49]. We mapped genes between mouse and human Entrez identifiers using vertebrate homology information from MGI, (http://www.informatics.jax.org/homology.shtml, downloaded March 2019). Module preservation calculations [50] were carried out using the modulePreservation function in WGCNA with 500 permutations. We summarized the calculated preservation statistics using the Zsummary measure.

## Data availability

Mice liver RNA-seq source data will be deposited at NCBI GEO dataset and will be publicly available. WGCNA original codes are publicly available at GitHub (https://github.com/plangfelder/Core-liver-homeostatic-networks). Any additional information required to reproduce this work is available from the Lead Contact.

## Authors’ contribution

Conceptualization, S.E.; Methodology, S.E., M.K.A, P.L., M.B., T.G.B; Project Administration: S.E., V.H., T.G.B; Resources: S.E., V.H., T.G.B, G.R.; Investigation, G.A.T, M.K.A., M.R-M, V.H., M.B., D.V., J.C., G.R., S.D., M.E., S.E.; Formal Analysis, P.L., D.V., C.L., B.S.G; Visualization, P.L., D.V.; Data Curation, S.E., P.L., T.G.B; Writing-Original draft, S.E.; Writing-Review & Editing, S.E., P.L., D.V., T.G.B, B.S.G, G.A.T, J.G.; Funding Acquisition; J.G., S.E.; Supervision, S.E, J.G..

## Acknowledgements

S.E. M.E. C.L. and J.G. are supported by the Robert W. Storr Bequest to the Sydney Medical Foundation, University of Sydney; National Health and Medical Research Council of Australia (NHMRC) Program (GNT1053206), Investigator and MRFF grants (GNT2032407; NCRI000183; GNT2016215; GNT2010795; GNT1196492) and a Cancer Institute, NSW grant (2021/ATRG2028). G.A.T. and D.V. are supported by a scholarship from the University of Sydney. Flow cytometry was performed in the Flow Cytometry Core Facilities that are supported by the Westmead Research Hub, the Westmead Institute for Medical Research, the Cancer Institute New South Wales, the National Health and Medical Research Council and the Ian Potter Foundation.

